# Chemical control and herbicide resistance of hairy fleabane [*Conyza bonariensis* (L.) Cronquist] in Jordan

**DOI:** 10.1101/2022.01.13.476249

**Authors:** Jamal R. Qasem

## Abstract

Two field experiments were conducted to evaluate the effectiveness of 12 herbicides in controlling hairy fleabane [*Conyza bonariensis* (L.) Cronquist] in a date palm orchard located in the central Jordan valley during the spring of 2017. Results showed that *C. bonariensis* resists paraquat (2.5, 5 and 7.5kgha^-1^), oxadiazon (5kgha^-1^) and oxyflourfen (3.3kgha^-1^) herbicides applied at normal or higher than the recommended rates. None of the three herbicides was significantly effective against the weed and treated plants continued growing normally similar to those of untreated control. Higher rates (10-fold of the recommended rates) of the same herbicides failed to control the weed. The effect of other tested herbicides on the weed was varied with bromoxynil plus MCPA (buctril^®^M), 2,4-D-iso-octyl ester, glyphosate, glyphosate trimesium and triclopyr were most effective and completely controlled the weed at recommended rates of application. Testing paraquat, oxadiazon and oxyflourfen using the normal recommended and 10-fold higher rates on two populations of *C. bonariensis* grown from seeds of the date palm and al-Twal (another site in the Jordan Valley) weed populations and grown in pots under glasshouse conditions showed that Date palm population was resistant to the three herbicides at both application rates while al-Twal site population was highly susceptible and completely controlled at normal and high rates of the three herbicides. It is concluded that certain populations of *C*. *bonariensis* developed resistance to paraquat, oxadiazon and oxyflourfen but control of this weed was possible using other herbicides of different mechanism of action. Herbicide rotation or other nonchemical weed control methods have been suggested to prevent or reduce the buildup and spread of resistant populations of this weed species. These results represent the first report on herbicide resistance of *C. bonariensis* in Jordan.

## Introduction

Herbicides represent the advanced weed control technology that utilizes synthetic and some bio-herbicides that cause inhibition or death of treated plants. The role of herbicides on the world food production is well known since their discovery in med. of the last century until now. However, this technology faces radical changes in effectiveness against a large number of weed species and biotypes under field conditions and lead in many cases to failure weed control due to continued development of weed tolerance/resistance and evolution and limitations in the herbicide industry and development [1]. Resistance means the survival and reproduction of herbicide treated plant after exposed to a rate of herbicide that is normally lethal to wild type [1, 2, 3, 4]. Therefore, it is a reduced response of a weed population to herbicides as a result of their selection pressure. However, when the same herbicide or herbicides of the same mechanism of action are repeatedly applied, resistant individuals increase in number and growth until dominate the site while sensitive individuals are suppressed or disappear. This however, takes a relatively long period for population shift from susceptible to totally resistant and depends on herbicide, environment and plant factors [1]. The resistance problem is becoming more severe in absence of other control methods or herbicide rotation [5]. In addition, the repeated use of low rates and/or applications beyond the label recommended growth stages and lack of tillage or other methods of control are also important contributing factors.

The genus *Conyza* includes 50–80 species spread in temperate and sub-tropical regions of America. Almost 50 of these species are spreading now as weeds in more than 40 crops in 70 countries [6, 7]. These have a high fitness and extremely efficient successful in spreading herbicide resistance within their population due to a combination of ecological properties such as a huge seed output, wide range of pollinating insects or self-fertilization, ability to outcross, short period for seeding, non-specific habitat requirements and ease and long distance seed dispersal. *C. canadensis, C. bonariensis*, and *C. sumatrensis* are most common while weed species include both diploids and polyploids [8] and can hybridize with each other [9].

*Conyza bonariensis* has been reported from different countries to resist glayphosate [10, 11, 12, 13, 14, 15, 16] and paraquat [17, 18] herbicides. The glyphosate rates required to control resistant populations were 7–10 times higher than those needed to control susceptible populations in southern Spain [19]. In addition, some biotypes have been mentioned to posses herbicide resistance in different parts of the world [3, 20, 21] and variations in resistance of the weed to both herbicides have been reported [22]. *Conyza canadiensis* has been also reported to resist glyphosate [5, 23], ALS inhibitors [24] and paraquat [18].

In Jordan, four *Conyza* species occur in the country but *C. bonsriensis* is the most common and widely spread in different regions in irrigated vegetables and fruit trees, waste places in the Jordan valley and high lands. It is an annual of 1-2 m in height, belongs to compositae family, and produces a huge number of modified hairy seeds that disperse far from parent plants. It flowers from April to November under local conditions.

In the last few years *C. bonsriensis* became more abundant, invading new lands and appeared dominating certain cultivated fields in the Jordan valley. However, specific studies on its control or reports on its resistance to herbicides are lacking from the country. The weed has been observed occupying date palm orchards and taking over the entire fields in certain locations in the Jordan valley. Local farmers claimed that it is not readily controlled by certain commonly used contact herbicides such as paraquat. Therefore, it was thought important trying some available foliage applied herbicides against this weed species.

## Materials and Methods

Two experiments were carried out under field conditions in the central Jordan valley and a single pot experiment under glasshouses conditions at the University of Jordan. These were as follows:

### Field experiments

#### Experiment 1

The experiment was conducted on April 18, 2017 in a date palm orchard at Dahrat Al-Ramil, in the central Jordan valley. The area is located at 255 m below the sea level and characterized by its tropical climate. The soil is a sandy loam of 50% sand, 25% silt and 25% clay with 1.3% organic matter content and a pH of 7.8. Date palm trees were 15 year-old and the weed was spread almost over the entire field and growing in pure stand except in some few spot in which *Cynodon dactylon* and *Prosopis farcta* were found. *Conyza bonariensis* was at a density of 30 plants m^-2^ and plants were at full vegetative or pre-flowering growth stages.

A weed-infested piece of land was chosen for the study, divided into 45 plots each of 2*2 m. Twelve herbicides were used and applied in 14 treatments and another untreated three plots were included as control (Table 1). All herbicides are available in various forms in local markets and included 2,4-D-iso-octyl ester [62% (v/v); Esterdefore; Veterinary and Agricultural Products Mfg. Co. Ltd., Amman, Jordan]; Mecoprop [60% (v/v); CMPP; Atlas Interlates Ltd.]; Glyphosate [48% (v/v); Roundup; Monsanto Agriculture Company, Monsanto Europe S.A., Athens, Greece]; Glyphosate trimesium [38% (w/v); Touchdown; Syngenta, Syngenta Crop Protection Pty. Ltd., Macquarie Park, NSW, Australia]; Linuron + Terbutryn [Tempo 13.9% EC, Pubchem]; Triclopyr [61.6% (v/v); Garlon 4E; Dow Chemical Company LLC, Indianapolis, IN, USA]; Oxyfluorfen [24% (v/v); Goal; Rohm and Haas, Mozzata, Italy]; Ioxynil octanoate [25% (v/v); Hocks; Veterinary and Agricultural Products Mfg. Co. Ltd. Amman, Jordan]; Paraquat [20% (v/v), Gramaxone, Sandoz, UK]; Oxadiazon [25% (v/v); Ronstar, Bayer, Germany]; Bentazon [48% w/v,_Basagran, BASF, Germany] and a 20% (v/v) Bromoxynil plus 20% (v/v) MCPA mixture [Buctril^®^M; May and Baker Agrochemicals (New Zealand) Ltd., Wingate, Lower Hutt, New Zealand].

**Table 1.**
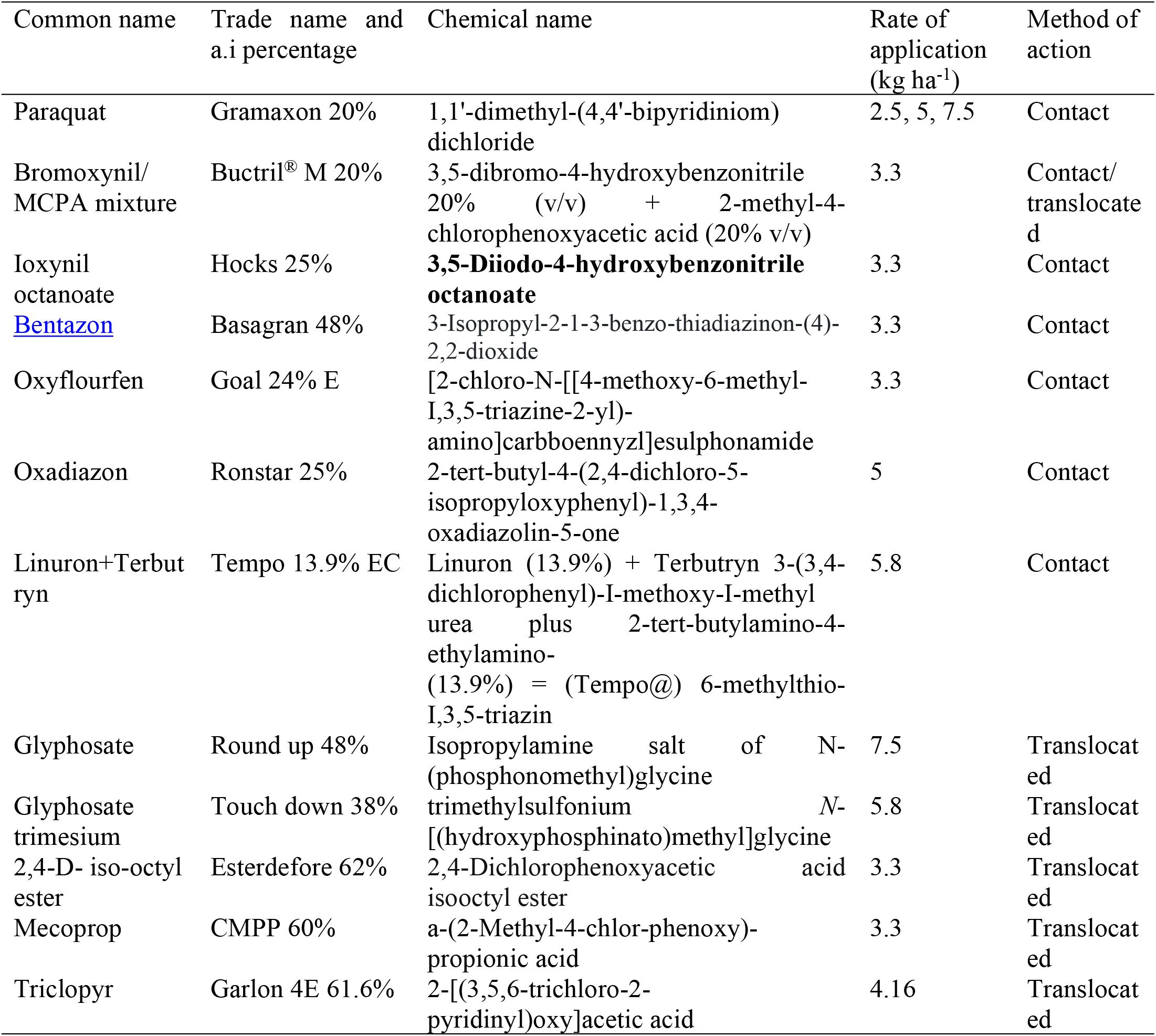
Herbicides tested for *Conyza bonariensis* control in the Date Palm orchard in the Jordan Valley in 2017.

All herbicides were applied as an aqueous spray on the foliage parts of the full vegetative to pre-flowering weed and at a constant pressure in the morning in absence of air movement, using a Knapsack sprayer with a single nozzle at a volume rate of 1083.3 l ha-^1^. All herbicides were applied on April 18, 2017 and visual estimation of the effects of these chemicals on weed growth was carried out twice on May 3^rd^ and 11^th^, 2017 using a scale ranged between zero to 10, at which 0 denotes that *C. bonariensis* plants were completely controlled and not recovered after treatment., and 10 means that no phytotoxicity effect of the herbicide was observed and the weed was in full growth. The evaluation was carried out by three persons and the average of the three scores given was considered for each plot. All plots were photographed and representative photos for severely affected or completely controlled *C. bonariences* plants and least or not affected (resistant) plants by certain herbicide treatments were included.

#### Experiment 2

The same procedure was followed as above in experiment 1, but only paraquat, oxyflourfen and oxadiazon herbicides were re-tested since the weed showed no response or was not affected by these chemicals at normal rates (and higher rates for paraquat) of application in the first experiment. The three herbicides were applied using10-fold the recommended rates on 18 May 2017 and similar scale of evaluation and estimation of the herbicides effects on the weed were carried out two weeks after application. Treatments replications and the layout of experimental design were the same as for experiment 1.

### Glasshouse Experiment

Seeds of *C. bonariensis* were collected from mature plants of the date palm orchard weed population in the central Jordan valley and al-Twal north almost 40 Km far from the first site but in the same region. Seeds of both populations were sown similarly in 10 cm diameter plastic pots filled with soil/peat mixture (1:1 V/V) at enough density on January 15, 2019. After emergences other weeds in all pots were hand removed and only *Conyza* plants were left growing. Plants of both populations were grown and irrigated as needed under unconditioned glasshouse. At full vegetative growth stage weed plants in each population (total 40 pots) were divided into two groups each of 20 pots that split into four subgroups each of five pots. Three sub-groups of five pots each were treated with paraquat, oxadiazon and oxyflourfen herbicides used at the normal recommended rates (2.5, 5 and 3.3kg^-1^ for each, respectively). Similar sub-groups were treated with the same herbicides but using 10-fold of the normal recommended rates of applications. Herbicides were applied using a PVC shoulder-held sprayer each contains 285 ml solution calculated to cover an area of 1m^2^ over which five pots of each population was spread over and treated with a single rate of one herbicide. The effect of herbicides on plants growth was visually estimated at five days after application using the same evaluation scale used for evaluation of the herbicides effects in field experiments. The same herbicides were re-applied on the same plants two weeks after the first application using the same rates used in the first spray. The effect of herbicides was visually estimated at five days after application using the same evaluation scale used before. In this experiment, each herbicide was sprayed twice and exactly as done in field experiments. However, since all herbicides tested are of contact action, complete coverage treatment was performed on both weed populations. Untreated pots (five pots of the fourth sub-group) were included as controls and for each weed population. Visual observations and herbicide effects estimates were noted and performed on all herbicides treatments five days after each application.

### Statistics

Treatments in field and glasshouse experiments were laid out in a randomized complete block design with 3 replicates for treatments in field experiments and 5 replicates for the second. Visual estimation scores on herbicides effects on weed populations and in all experiments were subjected to the analysis of variance (ANOVA) using SAS software SAS (r) version 9.1 [25]. Treatments means were separated and compared using the least significant difference test (LSD *p* = 0.05).

## Results

### Field experiments

#### Experiment 1

The effects of tested herbicides on *C. bonariensis* were visually estimated in the field at 2 and 3 weeks after treatments. Herbicides were greatly varied in their effects on the weed (Table 2).

**Table 2.** Visual estimations of the effects of foliage applied herbicides on *Conyza bonariensis* control, values are average scores of two estimations at 15 and 23 days after herbicides application (DAT).

| Herbicide | Score at 15 DAT | Score at 23 DAT |
| --- | --- | --- |
| Untreated (Control) | 10.0<sup>a</sup> | 10.0<sup>a</sup> |
| Paraquat (2.5 kg ha<sup>-1</sup>) | 10.0<sup>a</sup> | 10.0<sup>a</sup> |
| Paraquat (5 kg ha<sup>-1</sup>) | 10.0<sup>a</sup> | 10.0<sup>a</sup> |
| Paraquat (7.5 kg ha<sup>-1</sup>) | 9.0<sup>ab</sup> | 10.0<sup>a</sup> |
| Bromoxynil/MCPA mixture (Buctril® M) | 0.0<sup>e</sup> | 0.0<sup>e</sup> |
| Ioxynil octanoate | 3.0<sup>cd</sup> | 1.0<sup>d</sup> |
| Bentazon | 3.8<sup>c</sup> | 3.0<sup>c</sup> |
| Oxyflourfen | 9.8<sup>a</sup> | 10.0<sup>a</sup> |
| Oxadiazon | 7.8<sup>b</sup> | 9.5<sup>a</sup> |
| Linuron+terbutryn | 7.8<sup>b</sup> | 5.5<sup>b</sup> |
| Gluphosate | 1.8<sup>de</sup> | 0.0<sup>e</sup> |
| Glyphosate trimesium | 1.8<sup>de</sup> | 1.0<sup>d</sup> |
| 2,4-D- iso-octyl ester | 0.8<sup>e</sup> | 0.3<sup>de</sup> |
| Mecoprop | 7.3<sup>b</sup> | 2.8<sup>c</sup> |
| Triclopyr | 1.0<sup>e</sup> | 0.0<sup>e</sup> |
| LSD ≤ $p$ (0.05) | 1.8 | 1.0 |

Bromoxynil/MCPA mixture (Buctril^®^M) was best in controlling the weed and totally controlled all plants in treated plots shortly after application. This herbicide however, was followed by other highly effective herbicides including 2,4-D-iso-octyl ester, triclopyr, glyphosate and glyphosate trimesium. The ioxynil octanoate and bentazon showed moderate effects while paraquat, oxadiazon and oxyflourfen failed to affect the weed and to a less extent were linuron + terbutryn (Tempo*) and mecoprop herbicides.

Visual estimation of the effects of all herbicides treatments carried out at three weeks after application showed bromoxynil/MCPA mixture (Buctril^®^M), triclopyr and glyphosate had the highest effects and completely destroyed weed plants (Table 2 and Fig. 1) and these followed by 2,4-D-iso-octyl ester, glyphosate trimesium and ioxynil octanoate. The effect of mecoprop was much improved at this evaluation date but bentazon effect was slightly higher. However, the effect of paraquat used at increasing rates of application (2.5, 5 and 7.5 kgha^-1^), oxadiazon (5 kgha^-1^) and oxyflourfen (3.3kgha^-1^) remained the same with no obvious phytoxic effect on weed plants (Table 2 and Fig. 2).

**Fig 1.**
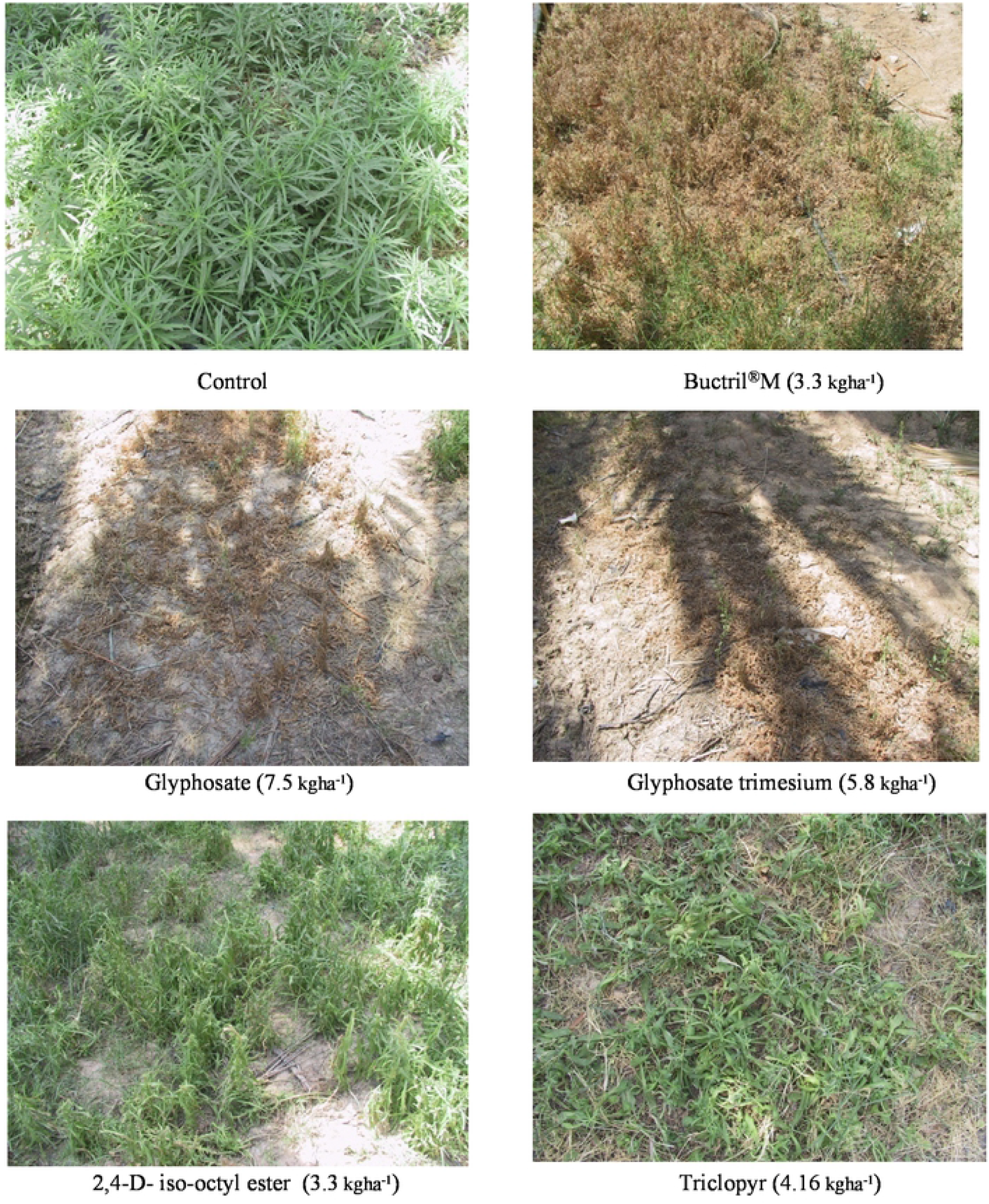
Most effective herbicides used at normal recommended rates of applications against *Conyza bonariensis* at 15 DAT.

**Fig. 2.**
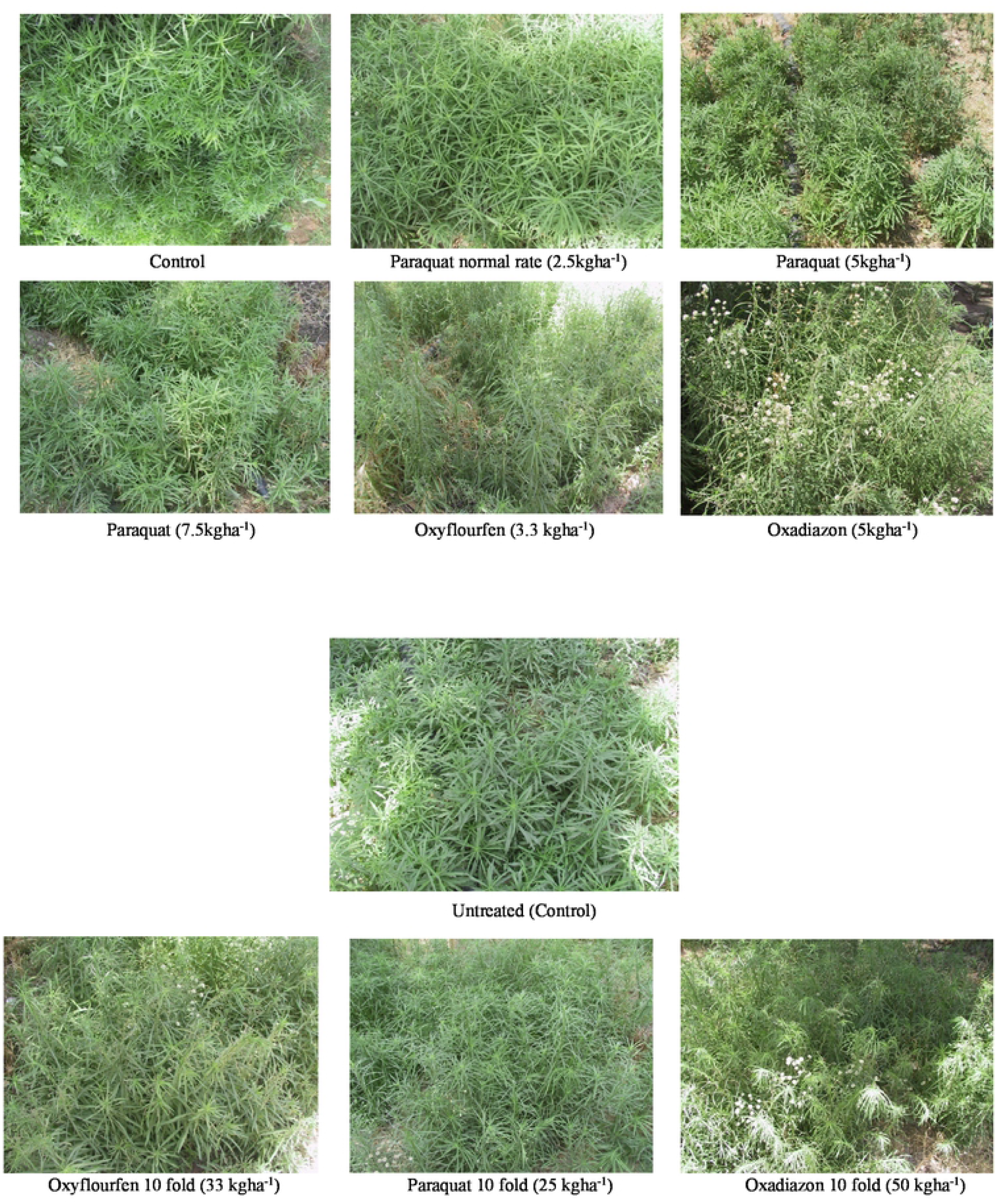
Effect of oxyflourfen, paraquat and oxadiazon used at normal and 10-fold of the recommended rat of application on *Conyza bonariensis* at 15 DAT.

The average effect of herbicides on *C. bonariensis* plants at the two estimation dates is shown in Fig. 3. It is clearly shown that paraquat (at all application rates), oxadiazon and oxyflourfen failed to control the weed. In contrast, bromoxynil/MCPA mixture (Buctril^®^M), glyphosate and triclopyr were most effective and totally killed weed plants (Fig. 1). These chemicals however, were followed by 2,4-D-iso-octyl ester had almost similar effect, ioxynil octanoate and bentazon and mecoprop showed moderate effects while linurone plus terbutrine formulation (Tempo*) was slightly better than the least effective or failure herbicides.

**Fig. 3.**
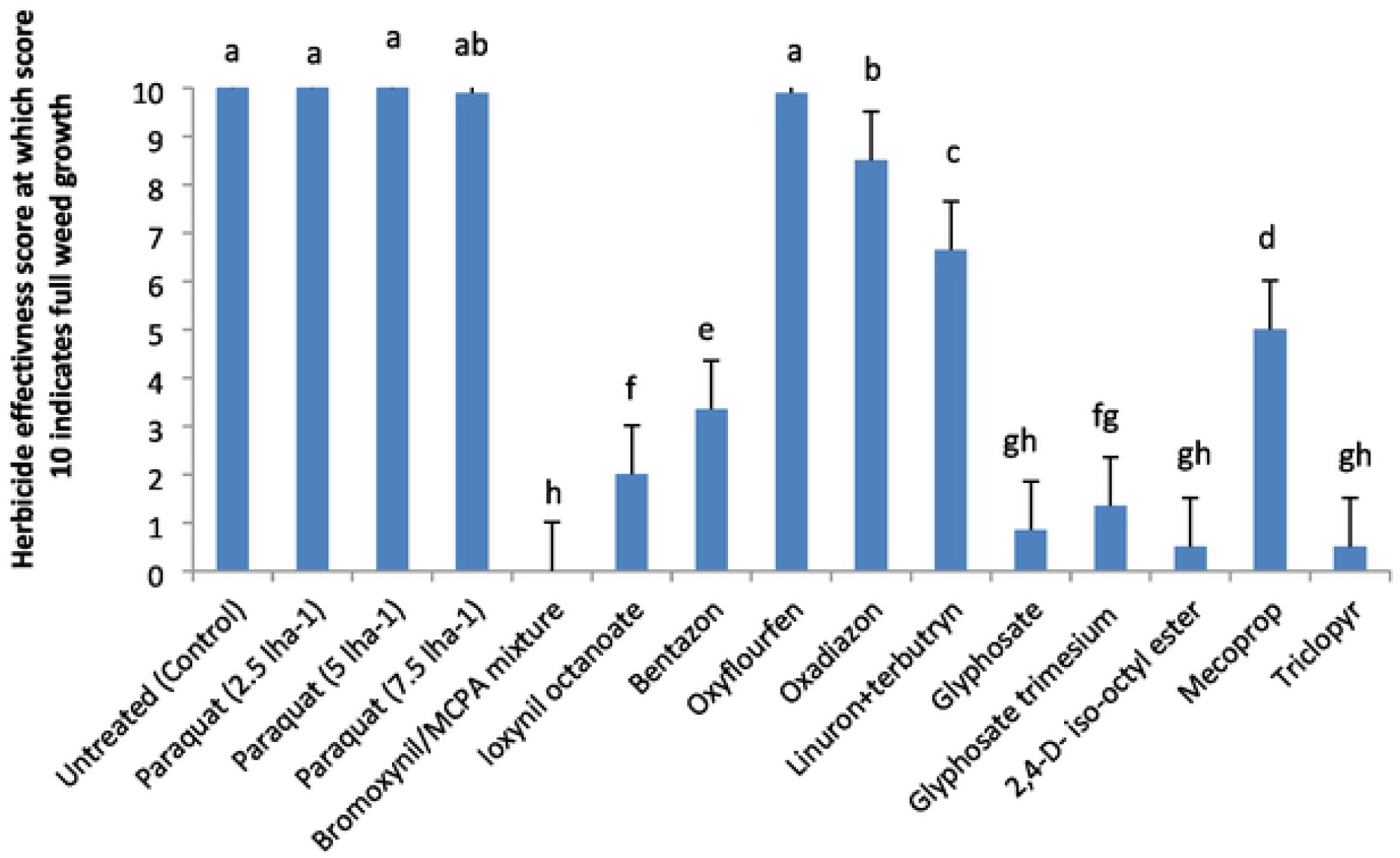
Average two estimates of the effect of different herbicides used for control of *C. bonariensis* in Date Palm orchard in the Jordan Valley. 1= LSD values at P ≤ 0.05. (columns represent growth of treated *Conyza bonariences* plants compared with untreated control). Mean values in the same column followed by the same lower-case letter arc not significantly different according to Fisher’s LSD at *P* = 0.05.

#### Experiment 2

Testing the highest rates (10-flod of the recommended rates) of paraquat, oxadiazon and oxyflourfen herbicides failed to control the weed (Table 3). At two weeks after application, none of the rates used for any of the three herbicides affected *C. bonariensis* growth but caused somehow a slight change in green color of the plants with some burning on leaf tips in oxadiazon and oxyflourfen treated plots (Fig.2). However, weed plants appeared normal and restored their green color at the second evaluation date three weeks after application (Table 3, Fig. 2).

**Table 3.** Visual estimations of the effects of selected foliage applied herbicides on *Conyza bonariensis* in date palm field, values are for two estimations conducted at 15 and 23 days after herbicides treatments (DAT).

| Herbicides | Rate of application<br>(kg ha<sup>-1</sup>) | Score at 15 DAT | Score at 23 DAT |
| --- | --- | --- | --- |
| Untreated (Control) | 0 | 10.0<sup>a</sup> | 10.0<sup>a</sup> |
| Paraquat | 25 | 8.0<sup>b</sup> | 9<sup>b</sup> |
| Oxyflourfen | 33 | 9.0<sup>ab</sup> | 10<sup>a</sup> |
| Oxadiazon | 50 | 9.0<sup>ab</sup> | 10<sup>a</sup> |
| LSD ≤ *p* (0.05) | - | 1.3 | 0.7 |

Mean values for five replicate pots treated with each herbicide on a scale of 0-10 where 0 denotes that the weed was completely controlled and the weed had no possible recovery after treatment., and 10 means that the weed was in full growth and no herbicide phytotoxicity observed. Mean values in the same column followed by the same lower-case letter are not significantly different according to Fisher's LSD at *P* = 0.05.

### Glasshouse experiment

The effects of paraquat, oxadiazon and oxyflourfen on *C. bonariensis* used at normal and high rates of application were shown in Table 4.

**Table 4.** The effects of low and high rates of application of three herbicides on two populations of *C. bonariences* grown from seeds collected from two sites in the Jordan valley and sown in pot experiment and sprayed twice with the herbicides at full vegetative growth stage.

| Treatments | Normal rates of application (kg ha<sup>-1</sup>) | First spray 15 October 2019<br>Score out of 10 | Second spray 30 October 2019<br>Score out of 10 | High rates of application (kg ha<sup>-1</sup>) | First spray 15 October 2019<br>Score out of 10 | Second spray 30 October 2019<br>Score out of 10 |
| --- | --- | --- | --- | --- | --- | --- |
| Date Palm Resistant Population of *C. bonariensis* |  |  |  |  |  |  |
| Control | 0 | 10.0<sup>a</sup> | 10.0<sup>a</sup> | 0 | 10.0<sup>a</sup> | 10.0<sup>a</sup> |
| Paraquat | 2.5 | 9.2<sup>a</sup> | 7.8<sup>b</sup> | 25 | 6.4<sup>d</sup> | 4.6<sup>c</sup> |
| Oxyflourfen | 3.0 | 9.2<sup>a</sup> | 8.4<sup>ab</sup> | 30 | 8.0<sup>bc</sup> | 7.4<sup>b</sup> |
| Oxadiazon | 5.0 | 9.4<sup>a</sup> | 8.4<sup>ab</sup> | 50 | 8.4<sup>ab</sup> | 7.2<sup>b</sup> |
| Al-Twal Susceptible Population of *C. bonariensis* |  |  |  |  |  |  |
| Control | 0 | 10.0<sup>a</sup> | 10.0<sup>a</sup> | 0 | 10.0<sup>a</sup> | 10.0<sup>a</sup> |
| Paraquat | 2.5 | 0.8<sup>c</sup> | 0.0<sup>d</sup> | 25 | 1.2<sup>f</sup> | 1.0<sup>d</sup> |
| Oxyflourfen | 3.0 | 0.6<sup>c</sup> | 0.0<sup>d</sup> | 30 | 3.4<sup>e</sup> | 1.0<sup>d</sup> |
| Oxadiazon | 5.0 | 8.2<sup>b</sup> | 2.4<sup>c</sup> | 50 | 7.2<sup>cd</sup> | 2.4<sup>d</sup> |
| LSD (*p*≤0.05) | - | 0.96 | 1.7 |  |  |  |

Mean values for five replicate pots treated with each herbicide on a scale of 0-10 where 0 denotes that the weed was completely controlled and the weed had no possible recovery after treatment., and 10 means that the weed was in full growth and showed no herbicide phytotoxicity. Mean values in the same column for both weed populations followed by the same lower-case letter are not significantly different according to Fisher's LSD at *P* = 0.05.

*Conyza bonariensis* was not affected by any of the herbicides used and at both rates of application. Weed plants were growing normally and more or less similar to those of untreated control (Table 4). In contrast, weed plants raised from al-Twal population seeds were completely controlled with paraquat and oxyflourfen but were less so using oxadiazon and at both low and high rates. However, repeated application of the same herbicides on weed plants of date palm population resulted in some more phytotoxic effects especially with paraquat but plants almost continued growing normally thereafter (Table 4). It was noted that herbicide-resistant plants were growing higher than herbicides-susceptible, unbranched or of low number of branches, late flowering and produced less number of flowers that consequently produced lower seed number compared with herbicides-susceptible plants (Fig.4).

**Fig. 4.**
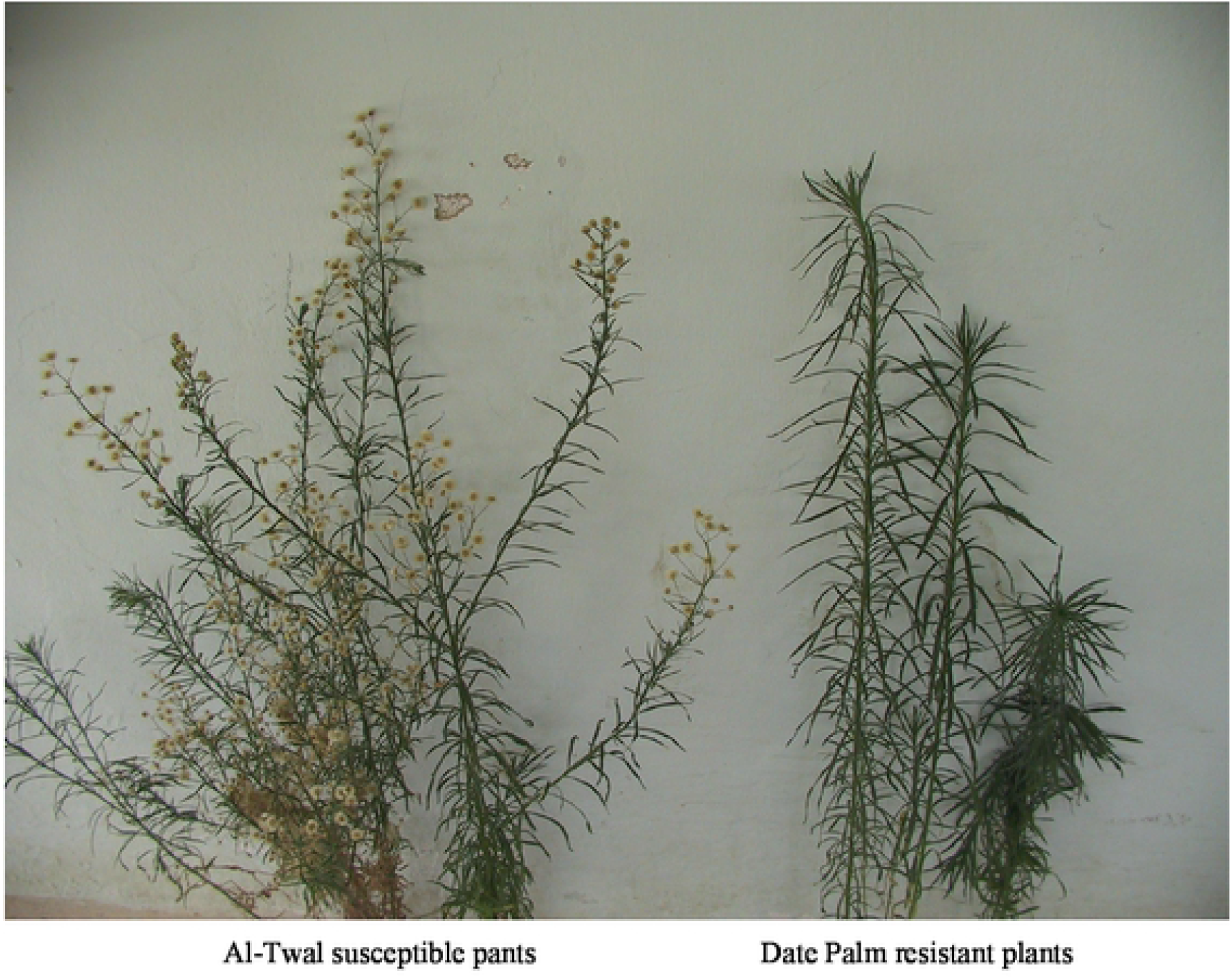
Herbicide-susceptible plant raised from seeds of al-Twal site population (left) and herbicide-resistant plant raised from seeds collected of the date palm orchard population (right) located in the Jordan valley.

## Discussion

Hairy fleabane [*Conyza bonariensis* (L.) Cronquist] populations in certain parts of the Jordan valley are rapidly expanded in the last few years. The weed invading vegetables as well as fruit tree orchards and can be observed in uncultivated lands and roadsides. It is also spread in the high lands in different field crops. Recently, some of the Date Palm farmers in the Jordan valley complained from that the weed is irresponsive to paraquat applications and is not readily controlled using this chemical. Praquat was widely used in the country for general weed control and in fruit tree orchards. Although the chemical was banded few years ago but stock of this chemical is available in stores of different agricultural companies and the herbicide is still marketed under other trade names. However, weed species resistance to this herbicide is expected since widely used by local farmers because cheap and rapidly desiccating weed plants shortly after application. Therefore, low price, low application rate and rapid effects on weeds all favored the herbicide by local farmers over other chemicals. This however, created serious ecological problems among which are paraquat-resistant weeds. Farmers however, trying other methods of weed control including tillage and weed grazing by cattle but both remain ineffective because of many weed characters among which the huge number of modified seeds produced and its ability to re-vegetate after grazed. Cattles partially graze this weed species and mostly feed on shoot tops and thus enhanced the development of auxiliary buds and branches that enable weed growth recovery and more seed production.

In the present study, beside paraquat, the widely used herbicide by farmers for general weed control in orchards, other herbicides were tested for possible control of paraquat-resistant *C. bonariensis.* These included contact and translocated chemicals among which was glyphosate that already reported [13, 15, 16, 18, 26] as resisted by the same weed species. However, all chemicals were applied at full vegetative to pre-flowering stage of the weed.

Our results were pronounced in which paraquat used at the three rates of application in the first field experiment failed to harm the weed. Our findings agreed with results of other workers reported the same weed species as resistant to paraquat [17, 18, 27]. Other contact herbicides tested mainly oxadiazon and oxyflourfen are commonly used by local farmers for weed control in vegetables (mainly onion and garlic) as well as in fruit trees and for general weed control in uncultivated lands in different parts of the country. This may have increased the possible build up of resistance and its development in certain common weed species or populations to these herbicides. However, testing the three chemicals against *C. bonariensis* population in the Date palm orchard showed that all were highly resisted by *C. bonariensis* although some other less used herbicides such as Linuron + terbutryn formulation (Tempo*) showed slight harmful effects on the weed. Results on oxyflourfen effect appeared not compatible with those of Travlos & Chachalis [28] who reported this herbicide as useful for managing resistant accessions of *C. bonariensis* in a mixture with glufosinate. The effect however, seems due to glufosinate.

In contrast, bromoxynil/MCPA mixture (Buctril^®^M), was most effective and completely controlled *C. bonariensis* plants shortly after application and treated plants failed to recover. This mixture is made from bromoxynil of a contact action mixed with MCPA with a systemic action and both are formulated in a mixture commercialized as buctril^®^M. MCPA has been reported as an excellent herbicide to control the same weed species at all leaf stages and all weed biotypes could be effectively controlled [29]. Glufosinate is a contact herbicide with some systemic action and has been reported as effective in controlling susceptible, glyphosate and paraquat-resistant populations of *C. bonariensis* (18). In addition, diquat, glufosinate, or glufosinate + oxyfluorfen controlled glyphosate-resistant or - susceptible *C. bonariensis* [28]. These herbicides, along with various integrated management strategies, have good potential to manage or slow the spread of glyphosate resistance in this weed species.

Our results demonstrated that translocated herbicides including glyphosate and glyphosate trimesium were highly effective and almost completely controlled the weed at 2^nd^ evaluation date. However, both showed complete weed control a week after first evaluated explaining the systemic action of these herbicides that has taken longer to translocate inside the plant and to show apparent phytotoxic effects on treated weed plants. Glyphoste has been reported as ineffective on *C. bonariensis* and the weed resists treatment with this herbicide [11, 12, 13, 15, 16, 18, 26]. The high effectiveness of the herbicide used in two forms against the same weed species in the present study may be because the chemical until the forbidden of paraquat was not widely used in the country but only against perennials that difficult to control by other herbicides or using other methods of weed control. Therefore resistance to this herbicide in the country may still not developed in this weed species. 2,4-D and triclopyr were both effective and almost completely destroyed weed plants at the normal recommended rates. However, these have taken longer than other herbicides to fully exert their harmful effects on the weed which is normally expected for translocated herbicides. These chemicals however, had their own specific mode of action observed at first evaluation time which included leaf malformation, twisting, and growth abnormality and curling. Some of these chemicals however, have been recommended by other workers [30] for use against the same weed species either alone or in mixture with glyphostae or picloram. Mixtures of glyphosate with saflufenacil and glyphosate with 2,4-D were reported as effective in controlling susceptible, glyphosate-resistant, and glyphosate-paraquat-resistant populations of *C. bonariensis* [18]. Glyphosate-resistant weed was effectively controlled using 2,4-D and dicamba [31] and the weed responded differently to both herbicides. It is worth mentioning that both MCPA and 2,4-D and trichlopyr are of related groups and no kind of resistance to any of these herbicides have been developed or reported in this weed species.

Further testing of paraquat, oxadiazon and oxyflourfen showed that resistance to these chemicals by *C. bonariensis* is well developed and very high application rates of the three herbicides had no appreciable harmful effects on the weed. All treated plants maintained normal growth similar to those of untreated control. Resistance of the weed to oxyflourfen and oxadiazon herbicides represents the first report on *C. bonariensis* resistance to these chemicals. The use of other herbicides of different mode of action is necessary to mitigate the spread of resistant population and glyphosate and phenoxy herbicides have an important role at this stage either separately, in combination or when integrated with other methods of weed control. However, plant, herbicide and environmental factors have also an important role in counteracting or worsening the weed resistance problem.

## Conclusions

It is concluded that adoption of integrated management of *Conyza* is essential to ensure that plants of this fast adapting genus will remain under control in arable and perennial crops. Farmers certainly are unfamiliar with *C. bonariensis* resistance to some commonly applied herbicides in different parts of the country. However, this is the first report on resistance of this weed species in Jordan and on weed resistance to herbicides in general. Since the three resisted herbicides in this study are widely used in the country and most likely included in weed control programs, therefore managements are necessary to be taken to prevent the development of resistant populations of this noxious weed species, species of the same genus and of other weed species. Agricultural practices should aim at preventing the spread of herbicide-resistant populations and reducing the buildup of resistance to the widely used herbicides. This however, can be only achieved through herbicide rotation and by using other weed control methods such as mechanical and cultural or at least minimizing the use of resisted herbicides by the weed in places where *C. bonariences* is prevailing and commonly spread. Label warnings should be also indicated on all products of the resisted chemicals by the weed and better management of the selection pressure resulted from the use of these herbicides. All possibly used chemicals against *C. bonariences* however, should be integrated with other weed control methods in order to prevent escalating populations’ resistance.

## Acknowledgements

The author thanks Mrs. Loai Al-Batsh and Madi Al-Abaddi for their technical assistance in the field.

## Conflicts of Interest

No conflicts of interest have been declared.

## Notes

### Competing Interest Statement

The authors have declared no competing interest.

